# Investigating termite nest thermodynamics using a quick-look method and the heat equation

**DOI:** 10.1101/161075

**Authors:** Rémi Gouttefarde, Richard Bon, Vincent Fourcassié, Patrick Arrufat, Ives Haifig, Christophe Baehr, Christian Jost

## Abstract

Termite mounds are often cited as an example of efficient thermoregulated structures. Nest thermal stability can be critical for insects that are particularly sensitive to heat and desiccation. Few studies have measured internal temperature of termite nests with respect to environmental parameters, especially in Neotropical species. In this study, we analyzed the thermal profiles of different parts of *Procornitermes araujoi* nests, a neotropical mound-building termite of the Brazilian *cerrado*. To read into our dataset we first used rasterization, a method that allows a quick-look assessment of time-series. Our results show that nest architecture efficiently buffers against environmental temperature fluctuations while at the same time maintaining a relatively high internal temperature in the core. This rather stable internal climate follows nevertheless the external temperature long-term averages. Using a novel numerical scheme, we further show that the heat transfer dynamics are well described by the classical heat equation, with an additional heat source whose origin is discussed.

## Introduction

Nests are supposed to protect animals from hostile environmental conditions [1]. In social insects, nests can be considered as microclimate regulation devices (for gas exchange, temperature or humidity) and have intrigued researchers for decades [2–6]. They are also an inspiring source for human engineering applications [7–10]. Nest thermoregulation can be characterized as either active or passive [4]. Active thermoregulation is defined as mastered by insect behavior such as fanning behavior in bees [4] or the creation of an energy sink by bringing up underground water in termites [11]. On the other hand, passive thermoregulation relies on mechanisms such as nest site selection [4,12] or nest architecture [6]. An outstanding example of passive thermoregulation is the ventilation system developed by some ant and termite species to increase the ventilation of their underground chambers [7,13–17].

Nest temperature is highly influenced by external periodic forcing. Even if the nest structure generally permits a good regulation of the internal microclimate, nest temperatures often follow short term (daily) and long term (weekly, annual) environmental fluctuations [7,18]. In mound-building species, like the African termite *Trinervitermes sp.,* the temperature fluctuations are more important in the upper part of the mound than inside the core [18,19]. Here, we investigate temperature dynamics in the Neotropical termite *Procornitermes araujoi* Emerson (1952) [20]. This species is endemic to the Brazilian *cerrado* (savanna). Its colonies inhabit nests that are characterized by medium-sized mounds usually rising less than 1m above ground (**Fig 1a**). The internal architecture is relatively simple with an homogeneous foam-like structure (**Fig 1a**) [21,22] also found in *Trinervitermes sp* or even in ant species like the black-garden ant *Lasius niger* [23]. It lacks the complex ventilation systems found in fungus-growing termites [15,24] or in leaf-cutting ants [14]. It is thus a convenient species to study nest thermoregulation in simple architectures with lower metabolic activity compared to fungus growing species.

**Fig 1.**
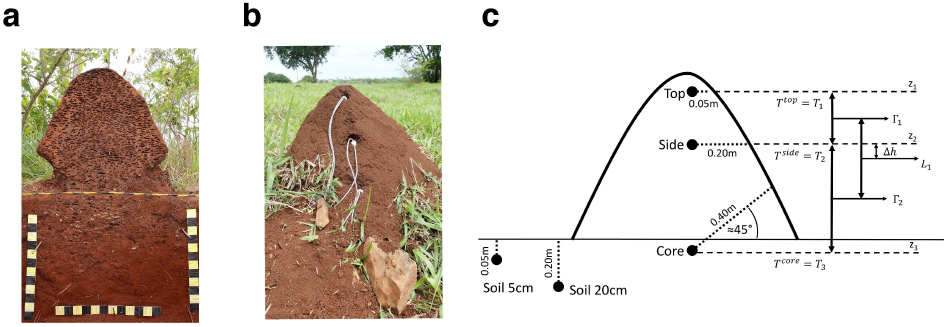
Experimental setup. (a) Sagittal section of a *P. araujoi* nest showing the foam-like structure and the underground extension of the nest. Yellow or black squares are 5 × 5 cm. (b) *P. araujoi* nest with the monitoring wires. (c) Probe positions inside and around the nests and spatial parameters used in the heat equation. T_1_ = Temperature of the top; T_2_ = Temperature of the side; T_3_ = Temperature of the core; Γ_1_ = heat flux between the top and the side probes; Γ _2_ = heat flux between the side and the core probes; Δ h = height difference between the time derivative estimation 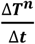 and the temperature heat flux gradient estimation 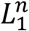 (see text for equations).

In closed mounds like those built by *P. araujoi*, nest material is likely to influence thermoregulation by affecting the microstructure of the walls and water retention, which are probably also crucial for gas exchange properties in such nests [16,25]. In *P. araujoi,* the workers use soil, regurgitated soil, fecal material and saliva as construction material [21]. Moreover, the mounds are often covered by a thin layer of loose soil and grass [20; personal observations] that could also have insulating properties [26].

In this study, we monitored nest temperature in different parts of *P. araujoi* nests. Our aim was to characterize the buffering effect of the structure, that is the ability of the structure to regulate the internal temperature against external environmental fluctuations. Previous studies have shown that mounds are subject to daily and long term temperature fluctuations and that termites may contribute to this buffering effect [11,18]. To investigate nest thermoregulation, we propose a methodology that first identifies thermal patterns and then to quantify them using the heat equation.

From a technical point of view, detecting patterns in raw time series (as our data) can quickly become tricky [27–29]. Such technical difficulties can obfuscate the actual biological interpretation of the data. Here we first propose to use a quick-look method: rasterization [30–32]. This method consists in transforming 1D time series with a dominant periodic forcing (here daily fluctuation) into a 2D image that can be suitable for a quick visual inspection. In addition we transformed the data by using different thresholding and normalization schemes that allows to accentuate patterns in the rasterized images and thus helping to understand the overall dynamic patterns. This rasterization method first allows us to detect the nests’ buffering efficiency at short (daily) and long (across days) scale. We then investigated the physics of the heat transfer dynamics inside the nests in the context of the heat equation. This equation predicts that (a) the mean temperature is independent of the monitored location in the nest, (b) the temperature amplitude decreases with increasing depth (distance to the top of the nest) *i.e.* shows mitigation, and (c) the phase shifts according to depth. A numerical scheme allows to fit the heat equation to the monitoring data. This fit provides two parameters to characterize the mound: the heat diffusion coefficient that quantifies the buffering effect (decreasing amplitude or mitigation), and a constant that tells whether the living mound is globally a free system, an energy source or an energy sink.

## Methods

### Biological material and experimental field area

Three nests of the termite *P. araujoi* were monitored during two weeks (04/11/2016- 18/11/2016) in a pasture located at the *Fazenda Capim Branco* (18°52’48’’S, 48°20’27’ ‘W), an area belonging to the Federal University of Uberlândia, Minas Gerais, Brazil (**S1 Fig**). The circumference of each nest at different heights (corresponding to the levels where each probe was inserted (**Figs 1b,c** and **Table S1**) was measured to estimate the size of the nests. The total volume of a nest was estimated as the sum of the volumes of each section. The volume of the upper section (top of the nest) was calculated using the formula for the volume of a cone (π × *R*^2^ × *h* / 3, with *h* the cone height and *R* the radius at its base). For the 2 lower sections, the formula of a truncated cone 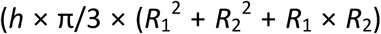 was used, where *R*_1_ and *R*_2_ are the radiuses of the upper and lower discs respectively). The total volumes were 0.34, 0.52 and 0.39 m^2^ respectively for nests A, B and C (**Table 1**). Taxonomic identification of the termite species was done using the soldier morphology following the dichotomous identification keys of Constantino [33] for the genus and Cancello [34,35] for species identification.

**Table 1.** Nest characteristics. Height H (m) and circumference C (m) of the 3 nests at different levels corresponding to the probe entrance points (see Fig 1c) with the resulting nest volume V.

| Nest | Top of the nest |  | Top probe entrance |  | Side probe entrance |  | Core probe entrance |  | Basis |  | V |
| --- | --- | --- | --- | --- | --- | --- | --- | --- | --- | --- | --- |
|  | H | C | H | C | H | C | H | C | H | C |  |
| A | 0.72 | 0 | 0.6 | 1.02 | 0.48 | 1.77 | 0.3 | 2.47 | 0 | 4.03 | **0.34** |
| B | 0.58 | 0 | 0.53 | 1.3 | 0.43 | 2.4 | 0.25 | 3.92 | 0 | 4.42 | **0.52** |
| C | 0.7 | 0 | 0.51 | 1.1 | 0.4 | 2.03 | 0.19 | 3.27 | 0 | 4.7 | **0.39** |

### Nest monitoring

Custom made devices (**S1 Text**) were used to measure the temperature every 10 min at five different positions, following Field & Duncan’s protocol [18] (see **Figs 1b,c** and **Table S1** for details). Each probe was placed into a 2 cm diameter plastic tube. The tubes were sealed at one end with a mesh to prevent the termites from entering. The other end was sealed with silicone. A concrete drill of 2.5 cm diameter was used to dig the holes into the nest (from the east side) and the ground where the tubes were placed. For each nest, two probes were placed inside each tube at each position in case of failure of one. Probes were placed (1) at the top of the nest, 5 cm below the nest surface, (2) on the side, at a horizontal distance of 20 cm from the surface, (3) inside the core of the nest (40 cm depth with an angle of 45° compared to the horizontal), in the soil next to the nest (4) at 5 cm depth and (5) at 20 cm depth, (6) at 1 m distance from the nest at 20 cm depth. When all the probes were installed, the data-loggers were started. The probes placed at the same location gave quasi identical data (**Table S1**). Therefore, only one of the two probes was used for the analysis.

Environmental parameters (air temperature, solar radiation, precipitations and relative humidity) were monitored in the meteorological station of the *Fazenda Capim Branco* situated approximately at 500m from the experimental field area.

### Data visualization: the rasterization method

Rasterization consists in transforming 1D time-series (see different examples of theoretical signals in **Fig 2a**) data into a 2D image by plotting the principal period (*e.g.* day, year, lunar month) on the y-axis, and subsequent periods on the x-axis; the actual signal is encoded in colors to obtain a heatmap like image (**Fig 2b**). Converting a time-series into a 2D image not only allows decomposing the signal, but also applies a smoothing effect that reduces the details and emphasizes the general patterns.

**Fig. 2.**
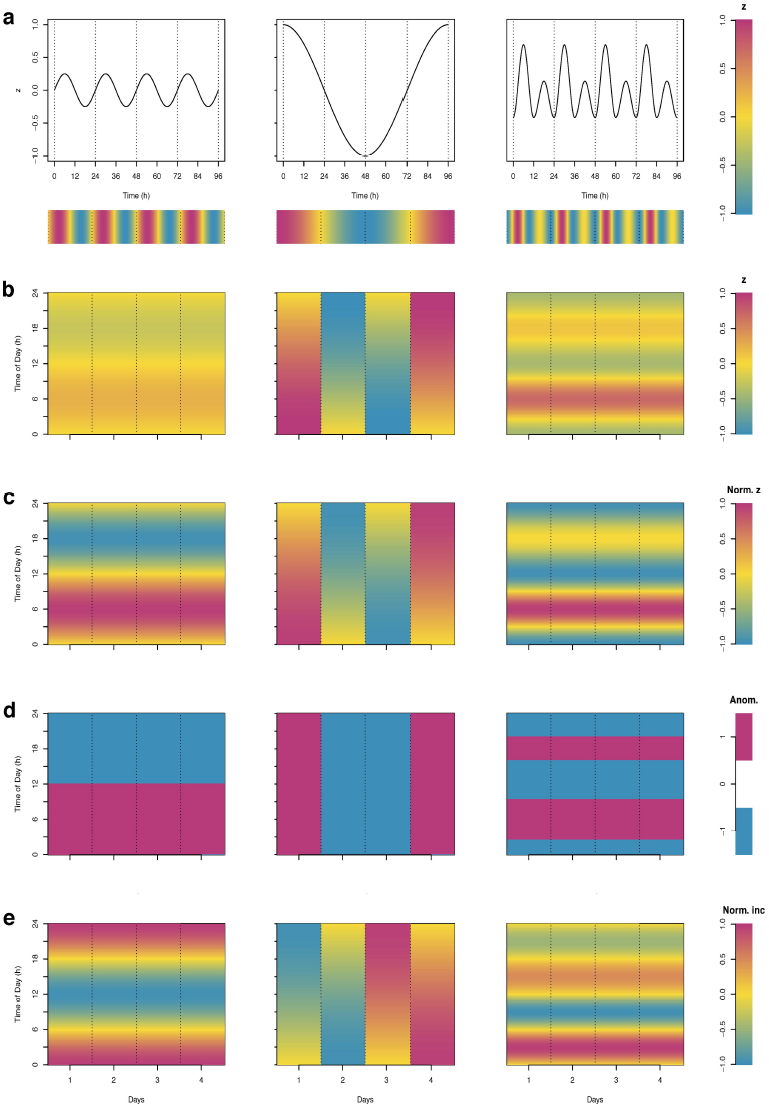
Illustration of the rasterization method. (a) “classical” 1D time-series for different kinds of theoretical signals sampled hourly. The horizontal color bar under the plots represents the encoding of the signal’s value in a color scale. Rasterized images of (b) the raw data of the 1D time series, (c) of the normalized data of the 1D time series, (d) of the sign of the departures from a reference value (anomaly), here the mean (values higher than the mean are plotted in magenta and those lower than the mean in blue), (e) of the normalized hourly increments.

In the case of a weak amplitude of the signal, some patterns can be masked by the color scale (**Fig 2b**). To solve this issue, one can enhance the contrast by normalization of the signal, for example into the range of [-1,1] (see **S2 Text eq (S2)**) (**Fig 2c**). This allows comparing easily the temporal dynamics of signals that have different amplitudes (**Fig 2a** and **2c**).

Another way to enhance contrast and facilitate pattern identification in rasterized images is to use hard thresholding (**Fig 2d**): it replaces each original value by one of two colors (binary data) according to a reference value used as a threshold. This allows to visualize the sign of the anomaly (that is defined as the departure of the value of a series from a reference value).

Finally, the dynamics of the time-series can be visualized by plotting the differences between subsequent values in the series (increments). These increments can also be normalized to increase contrast and facilitate pattern identification (**Fig 2e)**. All the plots were done with R 3.2.3 [36] (the dataset and an exemple of the script used to produce the figures are available online (http://doi.org/10.5281/zenodo.822263))

### Dynamical characterization using the heat equation

Here we investigate whether the spatio-temporal temperature evolution in the mound can be explained by a simple physical equation such as the heat equation. Using the *in-situ* temperature measurements, a numerical scheme of the heat equation is established in order to check whether this equation is able to predict the temperature inside the nest. This heat equation links linearly the temperature time derivative with the gradient of the heat fluxes. Both terms can be estimated from nest monitoring data if there are at least 3 probes.

### Mathematical background

The heat transfer equation describes the propagation of heat inside a domain. In the context of a termite nest heated by the sun, the heat transfer follows the sunray line. We assume that the heat propagation is done along the vertical axis and therefore that the domain is one-dimensional and semi-bounded. The equation describes the time dynamics of the temperature, represented by the time derivative of the temperature subject to the spatial variation of the vertical heat fluxes with a proportionality coefficient called the diffusivity coefficient.

The heat transfer is thus the partial differential equation

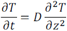

that may be also written as

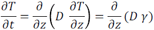

where *T* is the temperature (K) as a function of time *t* and depth *z, T* = *T t, z*, and 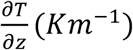 is the vertical heat flux. The boundary conditions of the semi-open domain are given by the surface temperature *T*_0_ = *T*(*t*, 0). The heat transfer coefficient *D* is a characteristic of the medium. In the present form of the heat equation, the medium is supposed to be homogeneous and *D* (*m*^2^*s*^2^) is a constant.

The proportionality between the temperature time 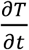 derivative and the heat flux gradient 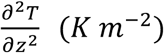 describes the flow of heat. It is well known that the solution of this equation is a diffusion process. In the case of a sinusoidal surface temperature, an analytical solution of the heat equation is known. We consider that *T*_0_ (*t*) = *T* (*t*, 0) = *A* sin (*ωt*) + *T*_*m*_where ω is the pulsation in *rad s*^-1^, *A* is the amplitude of the sinusoidal function and *T*_*m*_ is the mean temperature. Then the analytical solution is:

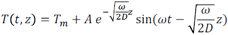

The solution of the above equation may be seen as the mean temperature, with an amplitude mitigation function relative to the depth and a phase difference in the sine function given by the depth. It is easy to transform the phase difference into a time lag. Indeed,

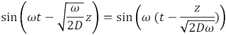

Therefore, the mathematical solution of the 1D heat transfer equation in the case of a sinusoidal forcing contains the two characteristics seen below in the nest temperature measurements: a time lag and mitigation according to the depth *z*. The heat transfer equation written previously corresponds to a free system. In the case of a non-autonomous system, with an energy well or forcing (such as heat absorption by water brought in by the termites or metabolic heat), the heat transfer equation is modified by an additive term,

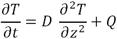

The additive energy *Q* is a negative term in case of a loss of energy, that is an energy sink, and non-negative for an input of energy, that is a positive forcing.

### Numerical approach

In order to link the heat equation to our temperature measurements at few discrete spots in the nest, it is necessary to discretize the partial differential equation.This discretization has to take into account the time characteristic of the cycles according to the time step of the data. It also has to take into account the heat propagation inside the soil. The flux transfer and the heat propagation in the soil are slow processes compared to external temperature forcing. Therefore, before the computation of the time and spatial derivatives, the fast variations are filtered from the temperature measurements by using a Fourier low-pass filter. Since the mean diffusivity coefficient and the mean additive forcing are of interest, the frequency of this low-pass filter corresponds to the diurnal cycle of 24 hours.

The time derivative is directly estimated by the first order difference: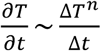 where Δ *t* is the data time step, and Δ *T* ^n^ is the difference Δ *T*^n^ = *T*^n^ - *T*^n-1^:; (*T*^n^ is the temperature at the time *t*^n^ = *n*Δ *t*). This temperature derivative series is estimated using the side probe *T*_2_.

The vertical temperature Laplacian at a given depth, Δ *T* is the temperature second derivative. This term is computed by two vertical derivatives. If 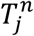 is the temperature for the time *t*^n^ = *n*Δ *t* at the j-th depth, the heat flux 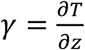 is estimated by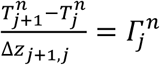 where Δ *z*_*j+1j*_ is the distance between the (j+1)-th and the j-th probes, *j* = 1 or 2. This flux is supposed to be estimated at the depth 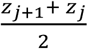. Using the three different depths, of our temperature measurements possible to compute two fluxes, and the second derivative of the temperature, which is the gradient of the heat flux, is then_computed with the same method in order to get 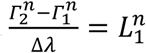 where 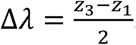. This term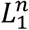 is supposed to be estimated at the depth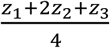

The estimates 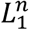 have to be corrected by a time shifting (associated to Δ *h* in Fig 1c) in order to be adjusted to the temperature increment. Indeed, the distances between the probes are quite large and the time lag is about one hundred minutes per 10 cm, whereas the time step is about 10 min. Therefore a time shifting of the Laplacian estimation *L*_;_ is necessary. Several methods are possible to assess this time shift (Fourier spectral shift, maximum of cross-correlation function, …). Here we estimated it as the number of time steps that maximize the absolute value of the correlation between temperature derivative 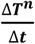 and the heat flux gradient 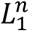

Finally, a linear regression between the time derivative 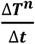 and the temperature second derivative 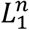 is performed. This regression provides estimates of the mean diffusivity coefficient *D* and of the mean additive forcing *Q*. Such a standard linear regression provides also the standard errors of these parameters. However, time series data do not fulfill the independence property assumed in these methods (our temperature time series are highly auto-correlated, leading to underestimated standard errors due to this pseudo-replication). Methods to correct for the bias induced by pseudo-replication are beginning to appear in the statistical literature [37] but their usefulness in the case of time series data is still strongly debated. We therefore chose not to report any standard errors for D and Q in a given nest, but to compute them with the classical formula over the three values obtained for our three (independently) monitored nests.

## Results

The mean, standard deviation and coefficients of variation for each probe are given in **Table 2.** The mean temperatures monitored at the different locations of the nest as well as in the soil are almost identical. For both the mound and the soil, the standard deviations decrease with depth z, leading to decreasing coefficients of variation. The temperature amplitudes are indeed much higher at the top of the nest (temperature range [16;44] °C) compared to the core (temperature range [24;28] °C; **Fig 3a, S2 Fig**). The same applies to the comparison of the soil temperatures at 5 and 20 cm. Mound temperatures are on average higher than soil and air temperatures (**Fig 3a**). Note that the air temperature and its amplitude are similar to those of the soil temperatures at 5 cm. In addition, daily variations of air temperature are intermediate between the top and the core of the nest.

**Fig 3.**
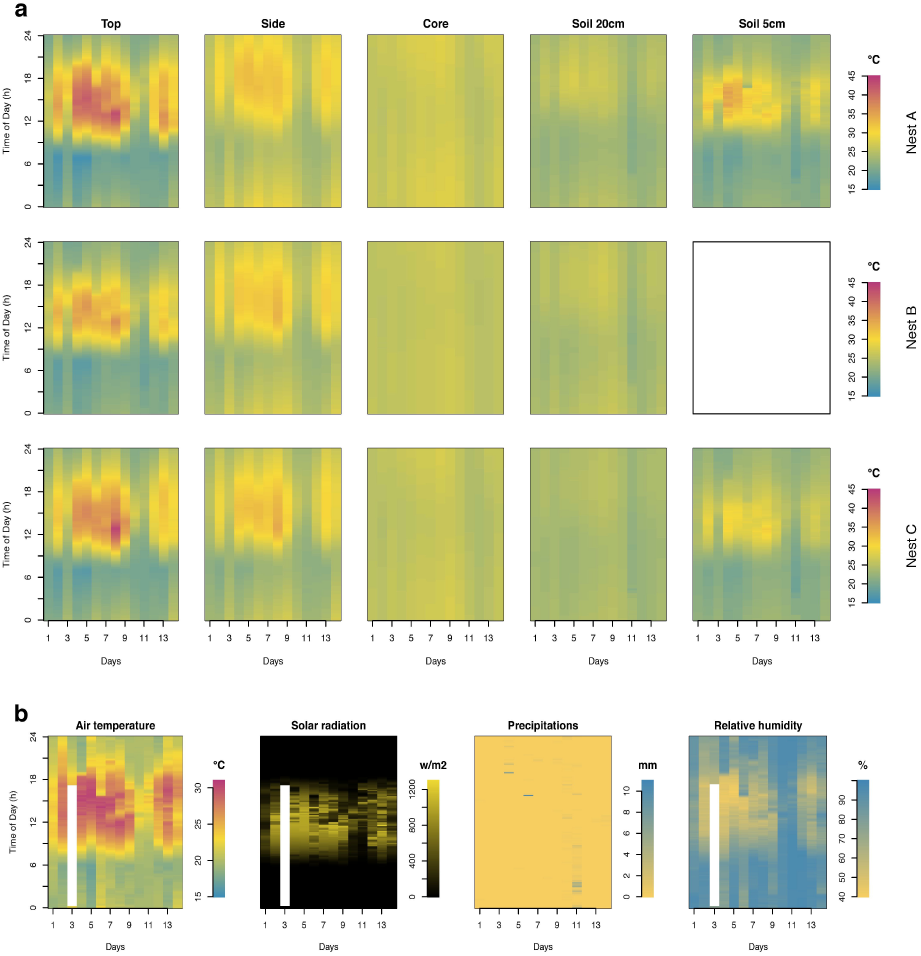
Rasterization raw data (transform 1D time-series into a 2D image). (a) for the 3 nests and all the probe positions and (b) for the environmental parameters (air temperature, solar radiation, precipitations and relative humidity). White space indicates missing data.

**Table 2.** Descriptives statistics. Mean (Mn), standard deviation (SD) and coefficient of variation (CV) of the temperature recorded by each probe for the 3 nests and for the air (meteorological station) during the 14 days. NA corresponds to the case where both inserted probes failed.

|  | Nest A |  | Nest B |  | Nest C |  |
| --- | --- | --- | --- | --- | --- | --- |
| | $Mn \pm SD$ (°C) | CV | $Mn \pm SD$ (°C) | CV | $Mn \pm SD$ (°C) | CV |
| Top | 25.82 ± 6.26 | 0.24 | 24.39 ± 4.46 | 0.18 | 25.54 ± 5.24 | 0.20 |
| Side | 27.00 ± 2.68 | 0.10 | 26.63 ± 2.73 | 0.10 | 25.89 ± 3.13 | 0.12 |
| Core | 26.42 ± 0.85 | 0.03 | 25.79 ± 0.52 | 0.02 | 25.64 ± 0.78 | 0.03 |
| Soil 5cm | 23.78 ± 3.42 | 0.14 | NA | NA | 23.59 ± 2.43 | 0.10 |
| Soil 20cm | 23.96 ± 1.34 | 0.06 | 24.50 ± 1.04 | 0.04 | 23.79 ± 0.72 | 0.03 |
| Air (station) | $Mn \pm SD$ (°C) | CV | | | | |
|  | 22.69 ± 3.36 | 0.15 |  |  |  |  |

### Interpretation of the rassterized images

The rasterized images of the raw data (**Fig 3a)** show comparable patterns for probes at the same location in the three nests. In addition to the decreasing amplitudes (mitigation)with increasing depth in both the nest and the soil, data also show that maximum temperatures are attained later in the evening with increasing depth (Fig 3a). However, on these figures one can barely see a pattern in the core or soil at depth 20cm. A clear pattern appears in both cases in **Fig 4** where we normalized the data into the range [-1;1]; maximum temperatures were much delayed compared to air and nest top ones (around 4 pm), occurring at midnight in the core. One can also see that the highest temperatures and the daily variation are smaller in all probe locations on days 11 and 12. This can be explained by cooler air temperature, less intense solar radiation and higher precipitations (with higher relative humidity, see **Fig 3b**) for these two days. This reveals that temperature of the nest follows the long-term environmental fluctuations and is true even for the core temperature although more attenuated than at the most peripheral locations in the nest and in the soil.

**Fig 4:**
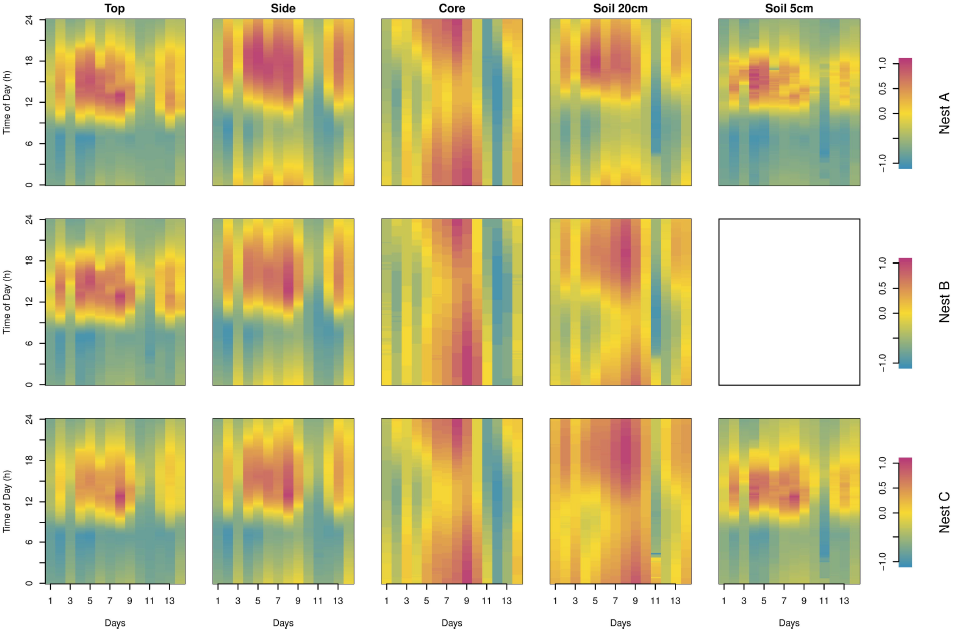
Normalized rasterized data. Data illustrated in Fig 3 (raw data) were normalized to restrict the values into a range of [-1;1] (see equation S1).

**Fig 5** shows the differential sensitivity of the upper part of the mound (top and side) and of the nest core to environmental fluctuations as revealed by the sign of the global anomaly of the time-series. The dominant vertical pattern seen in the core indicates that this latter is mainly sensitive to larger-scale external temperature changes while the dominant horizontal pattern seen in the top and side of the mound are more sensitive to the diurnal variation of the temperature. The pattern of anomaly shown in the soil at 5 cm and 20 cm depth are respectively quite similar to the ones observed at the top and the side of the nest showing that it follows principally daily fluctuations. The pattern of the side and soil at 20 cm reveals an intermediate pattern between the core and the surface (top of the mound and soil at 5 cm). Indeed, this pattern reflects both short (daily) and long (across days) term fluctuations. The relative long term fluctuations of the top and the side of the nest as well as of the soil at 5 cm can be visualized through the sign of the anomalies using the hourly mean temperature as threshold value (**Fig 6)**.

**Fig 5:**
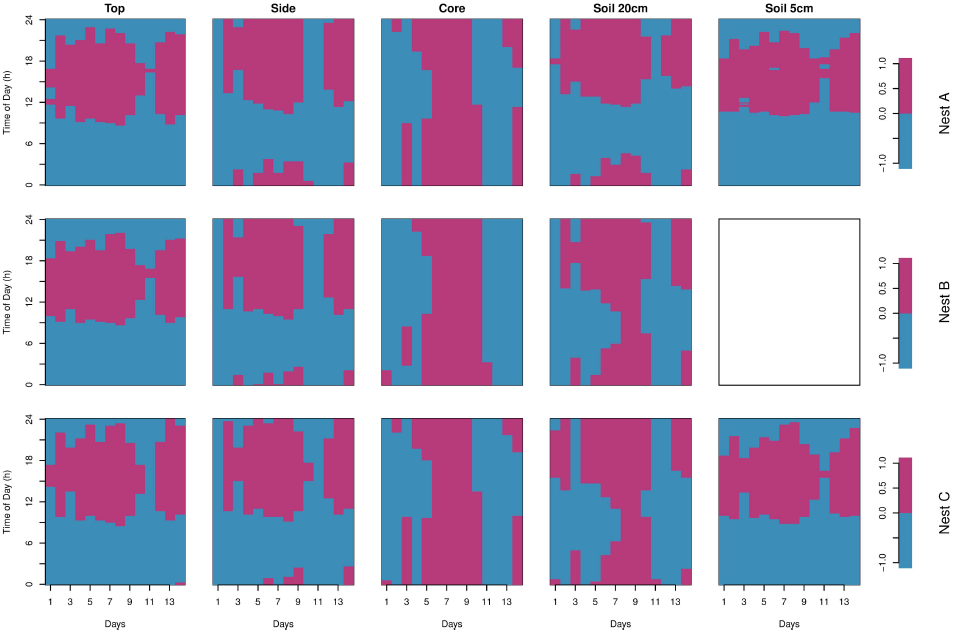
Rasterized sign of the anomalies using the overall mean as the threshold. The positive anomaly is indicated in magenta and the negative anomaly in blue.

**Fig 6:**
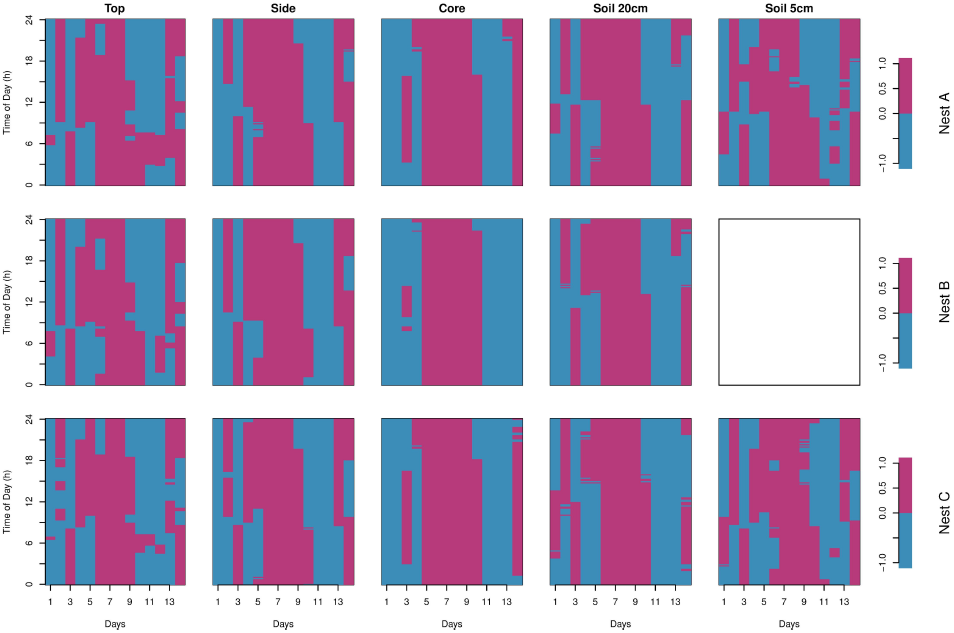
Rasterized sign of the anomaly using the hourly means over the two weeks of observation period as the reference thresholds. The positive anomaly is indicated in magenta and the negative anomaly in blue.

## Dynamical characterization

### Visualizing the heating and cooling phases

To visualize the temporal dynamics of heating and cooling, we plotted the hourly mean raw temperature at the different locations for the three nests (**Fig 7a**). We excluded days 11 and 12 which were different from the other days (**Fig 3**). The normalization of this data reveals a temporal shift of the maximum and minimum temperatures as one goes deeper inside the nest or in the soil (**Fig 7b**). Remarkably the heating and cooling of the core follows an inverted temporal pattern when compared to the top of the nest. Nevertheless, it is not easy to perceive whether the duration of the heating or the cooling are the same. This is easier to assess on the rasterized image of the normalized temperature increments following **eq S1** (**Fig 8**). The heating phase is well identified by the dominant magenta coloring, while the cooling phase has dominant blue coloring. Whatever the location, the heating phase is shorter than the cooling phase. We also notice that these two phases tended to be less asymmetric the deeper the probe location. The alternating pattern of 0 and of non-zero values in the least-varying core temperatures illustrates rather slow periods of heating and cooling (see **eq S1**). By contrast, the almost continuous color patterns at the top and the side of the nest indicate faster heating and cooling periods preceded or followed by a rather long period of stationary temperatures centered around midnight.

**Fig 7.**
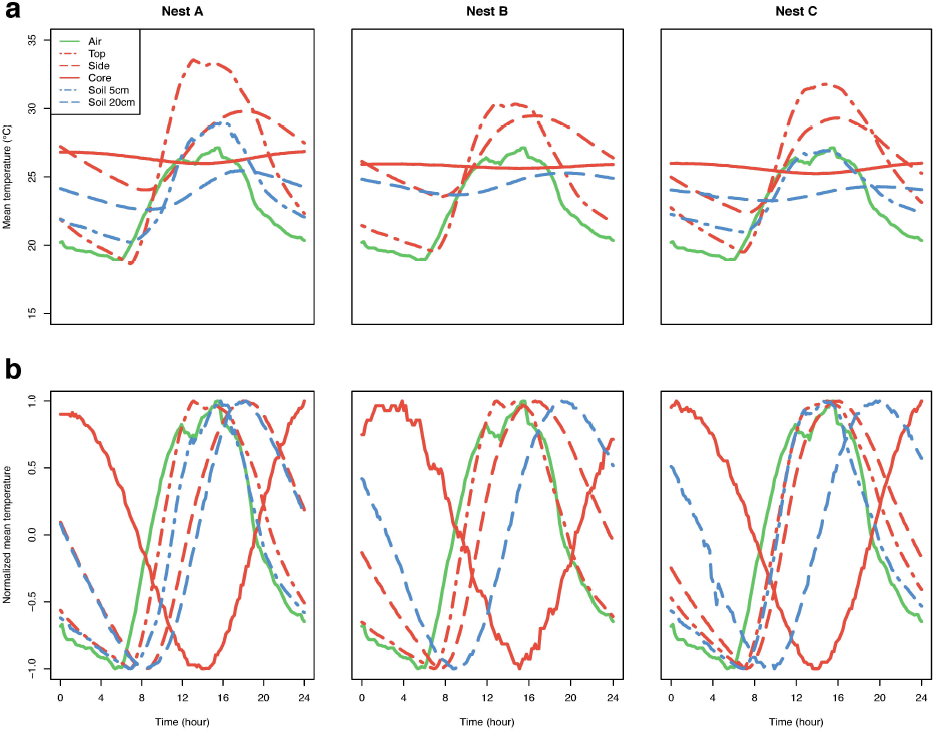
Mean temperatures along a 24h period. (a) Mean temperatures during the day after excluding days 11 and 12. (b) Normalized curves (between [-1 ;1]) of the same mean temperatures.

**Fig 8:**
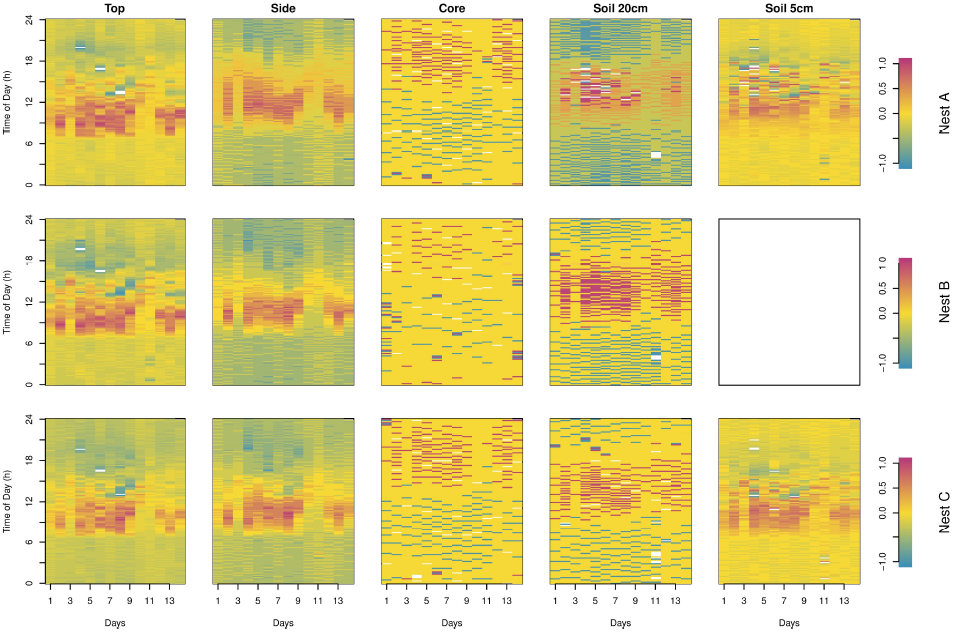
Rasterized increments of the raw data. The amplitudes of the increments Δ ***Tn*** are normalized to restrain the values into a range of [-1;1].

### Nest diffusivity coefficient

**Fig 7. Estimation of the temperature time derivative 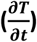 and the flux spatial gradient 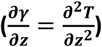 for nest A.** (a) time derivative as a function of the flux spatial gradient, with the best fitting line (red) to estimate the diffusivity coefficient ***D*** and the energy source ***Q***, (b) the time derivative and its linear prediction from the spatial flux gradient as a function of time, (c) the residuals of the linear regression in (a) as a function of time.

**Fig 9a** shows the relationship between the time derivative and the spatial flux derivative for nest A; the data are organized along a flat ellipse, a linear regression thus makes sense. The associated residual structure in **Fig 9c** confirms this conclusion. **Fig 9b** shows the time derivative and its linear prediction from the spatial flux gradient: there is a good agreement between the two series. The two other nests give quite similar plots.

The estimated parameters of the heat equation for the three nests are summarized in **Table 3.** Nests A and B have coefficients of the same order, with *D*~0.31 - 0.51 10^-6^*m*^2^*s*^-1^;. Nest C’s diffusivity coefficient is doubled. The mean additive forcings *Q* are all positive and in the range 0.37 - 1.24 10^-4^*Ks*^-1^. The time ag is about 70-80 minutes for nests A and B, and about 170 minutes for nest C. In conclusion, nest C seems to have a behavior different than the two other nests.

**Table 3.**
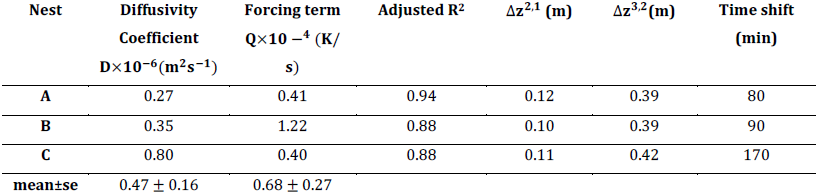
Estimated diffusivity coefficients and forcing terms in the three nests. The Δ *z* are the vertical distances between the probes and the time shift is the one used in the estimation of the heat flux gradients 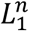

| Nest | Diffusivity<br>Coefficient<br>$D \times 10^{-6} (m^2 s^{-1})$ | Forcing term<br>$Q \times 10^{-4} (K/s)$ | Adjusted $R^2$ | $\Delta z^{2,1} (m)$ | $\Delta z^{3,2} (m)$ | Time shift<br>(min) |
| --- | --- | --- | --- | --- | --- | --- |
| A | 0.27 | 0.41 | 0.94 | 0.12 | 0.39 | 80 |
| B | 0.35 | 1.22 | 0.88 | 0.10 | 0.39 | 90 |
| C | 0.80 | 0.40 | 0.88 | 0.11 | 0.42 | 170 |
| mean±se | $0.47 \pm 0.16$ | $0.68 \pm 0.27$ | | | | |

## Discussion

Termite mounds are historically cited as an example of thermoregulated constructions [6]. Nevertheless, recent studies rather suggest that there is no active thermoregulation in these structures, even in the complex mounds of the fungus growing termite *Macrotermes michaelseni* [7]. Here, we measured temperature at different positions in the nest of the Neotropical mound-building termite *P. araujoi*. We aimed to characterize the dynamics of heat propagation and to investigate how the structure reacts to external forcing. We first used rasterized images to visualize our data and to decompose the effects of the diurnal and the large scale temporal forcing on temperature in the different parts of the nest. The results show that nest temperatures are strongly correlated with external forcing (solar radiation, rain), an argument for the weakness or absence of active thermoregulation. Nevertheless, even these simple homogeneous foam-like architectures show interesting thermal properties. Our results show indeed that the temperature pattern of the mound core differs from the top and the side of the mound as well as from the soil (both at 5 cm and 20 cm depth). On the one hand the core temperatures are very stable on a daily scale, while there are more important temperature fluctuations for the rest of the mound and the soil (**Fig 3** and **Table 2**). They are also higher than in the soil, thus providing a warm and very stable environment for the termites. On the other hand, it is also clear that the core, like the other parts of the mound and the soil, undergoes long term variations as the pattern observed for the core in Figs 4 and 5 corresponds to the one observed for the meteorological data (**Fig 3b**). These results confirm the findings on the mounds of the African termite *Trinervitermes sp*. which builds nests with similar architecture [18,19]. In particular, the increased core temperature has been found during the whole year [18] and is thus not only a simple consequence of monitoring temperature during the summer as we did.

From a functional point of view, the stable and warm temperature of the nest core might have significant effects on individual and colony development. Termites are hemimetabolous insects that undergo successive molting events during their life. In particular, during the post-embryonic development of termitid species, such as *P. araujoi*, the larvae follow one of either the apterous line, from which workers and soldiers originate, or the imaginal line from which alates originate [38]. Stable temperature may accelerate the rate of juvenile development [39], as it is the case in the nursery chambers of the nests of Macrotermitinae species (reviewed in [2]). Temperature can also influence caste composition [40,41]. Temperature variations may thus have a strong impact on the colony homeostasis which is based on the division of labor between the different categories of individuals which perform different behavioral activities [42].

In contrast to nests of *T. trinervoides* [18] our species’ nests extend below ground. A comparison between the core and soil temperature dynamics is therefore relevant. Though the patterns observed for the core are relatively similar to the ones for the soil at 20 cm depth, there are strong differences between the two concerning the dynamics of cooling and heating (**Figs 7** and **8**). The heating phase occurs later in the core compared to the soil. This could be explained by the fact that it takes much more time for the heat to propagate from the top of the mound which lies at 60-70 cm above the soil, than it takes to propagate from the soil surface to the 20 cm depth probe. Moreover, in the increments plot (**Fig 8**) one can see that the spacing between the discrete lines during the heating phase is tighter in the soil than in the nest’s core, showing clearly that the latter has more efficient buffering properties against external forcing. Compared to the soil, the mound core therefore provides a more stable (and warmer) environment for the termites. Note that temperature increments are negatively correlated with spatial temperature variation between probes (**Figs 4** and **8**), indicating that it is relevant to analyze nest temperature dynamics in the context of the general heat equation. In fact, our analysis suggests that the heat equation explains well the diffusion of heat inside the mound structure. Our estimation of the heat diffusivity coefficients are similar to those of soils of similar composition [43]. The higher diffusivity coefficient and time shift of nest C (Table 3) could be natural variation in our small sample or the particular position of this nest at the base of a steep slope in the field (Fig S1). To further investigate the effect of architecture or topographic environment on heat diffusion the mound and soil diffusivity should be compared in a paired design (at least 3 probes in both the nest and the adjacent soil) with more monitored nests. Bristow *et al* [11] had already used the heat equation to detect an energy sink in the mound-building termite *Tumulitermes pastinator*. Our numerical scheme goes significantly further in terms of model validation, precision and parameter estimation. The results obtained in *T. trinervoides* [18] suggest that mound temperature is increased in the presence of termites. However, simple metabolic heat has been ruled out in the case of *M. michaelseni* [7] which, as a fungus growing termite, produces much more metabolic heat than *P. araujoi*. An increase in temperature could be induced more actively by the termites [44] or by the mound design itself. Whatever the origin of this additional source of energy, our methodology permits to precisely quantify its value.

We think that the methodology we proposed here could be useful for future studies aiming to understand the mechanisms underlying termite nest thermoregulation. From a physical and evolutionary point of view a comparative approach between sympatric species that only differ by their mound architecture would be particularly interesting. In the case of *P. araujoi* we suggest to compare to *Cornitermes cumulans* (whose individuals are close in size but whose mound architecture is much more elaborated with solid outer walls and soft organic inner space [45,46]) or to *Cornitermes bequaerti* (whose mounds are not closed but have outside openings for ventilation [46,47]).

To conclude, rasterization allowed a quick assessment of temperature monitoring data, suggesting the heat equation could govern nest temperature dynamics. The parameters of the heat equation were estimated from the monitoring data in order to characterize the overall nest temperature dynamics. To our knowledge this is the first study that develops a numerical scheme to link the heat equation to mound temperature dynamics and thus validates its pertinence for the studied system.

## Acknowledgements

We would like to thank Tiago F. Carrijo from the Universidade Federal do ABC for his help with termite taxonomy and field work and Felipe Magalhães from the Universidade Federal de Uberlândia for providing the experimental field area. RG and VF received a financial support from the Université Toulouse 3 – Paul Sabatier for their field trip in Brazil.

## Author contributions

RG, RB, VF and CJ designed the experiments. RG, VF and IH did the field work. IH provided field support. PA designed and constructed the data logger. RG, CB and CJ did the data analyzes and wrote the manuscript. All authors discussed and commented on the manuscript.

## Supporting Information

**S1 Text. Data loggers.** Data loggers are designed to work autonomously for several weeks. They store temperature every ten minutes on an SD card (Secure Digital). The data logger has 3 main parts:

– 10 sensors measuring temperature (SHT25, Sensirion AG, Staefa ZH, Switzerland)
– Mass storage (2GB micro SD card, Transcend Information - Inc., Taipei, Taiwan)
– Microprocessor (PIC18F26K22, Microchip Technology, Chandler, Arizona, United States)

The processor coordinates the actions via an internal clock activated every ten minutes. It sequentially accesses each of the ten sensors, reads the temperature and stores these ten values in an internal RAM (Random Access Memory). Every 40 minutes, data in RAM are converted to ASCII (American Standard Code for Information Interchange) and then saved on the SD card in a standard CSV format (Comma-Separated Values). This implementation contributes to the long autonomy of the device. The data is coded on sixteen bits (maximum 65535 or five ASCII characters). The fourteen most significant bits represent the measured value and the two least significant bits contain status information that is not used here. These two bits must be set to zero for the calculation of temperature detailed below. Temperature T (*S*_*t*_ in binary format) is obtained through the transformation:

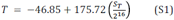

According to the SHT25 datasheet, the accuracy tolerance of the sensor is +/- 0.2°C in the normal range of use. For further energy economy the resolution of the analogue-to-digital conversion was chosen on 11 bits (faster reading times), which allows a resolution of approximately 0.08 °C (linearized values).

**S1 Table.**
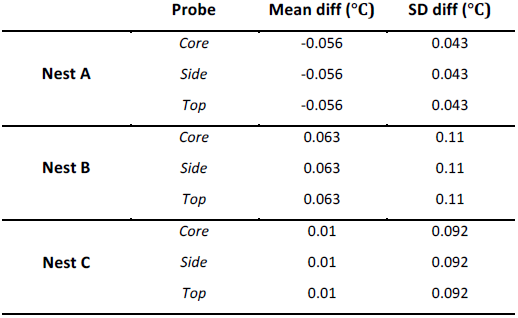
Differences between homologous probes. For all the nest positions, two probes were placed in the same tube. We calculated the mean and standard deviation of the differences for each of these pair of probes for the 3 nests.

|  | Probe | Mean diff (°C) | SD diff (°C) |
| --- | --- | --- | --- |
| Nest A | Core | -0.056 | 0.043 |
|  | Side | -0.056 | 0.043 |
|  | Top | -0.056 | 0.043 |
| Nest B | Core | 0.063 | 0.11 |
|  | Side | 0.063 | 0.11 |
|  | Top | 0.063 | 0.11 |
| Nest C | Core | 0.01 | 0.092 |
|  | Side | 0.01 | 0.092 |
|  | Top | 0.01 | 0.092 |

**S2 Text. Data normalization.** The data presented in **Figs 1e**, **1f** and **1i** were normalised using the following formula:

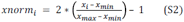

After applying this transformation, the new data values *xnorm*_*i*_ are bound between -1 and 1.

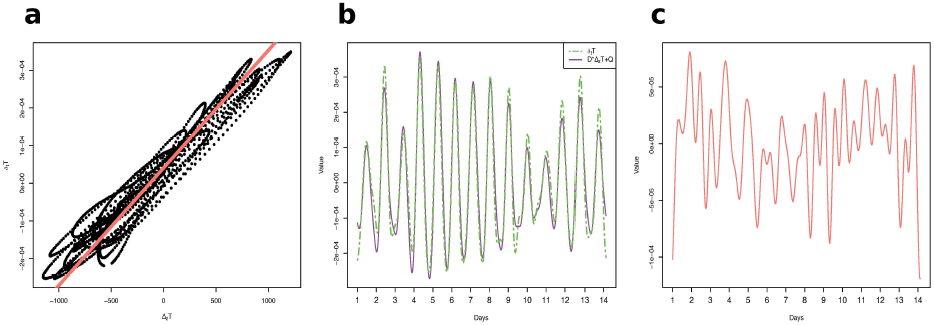

**S1 Fig.**
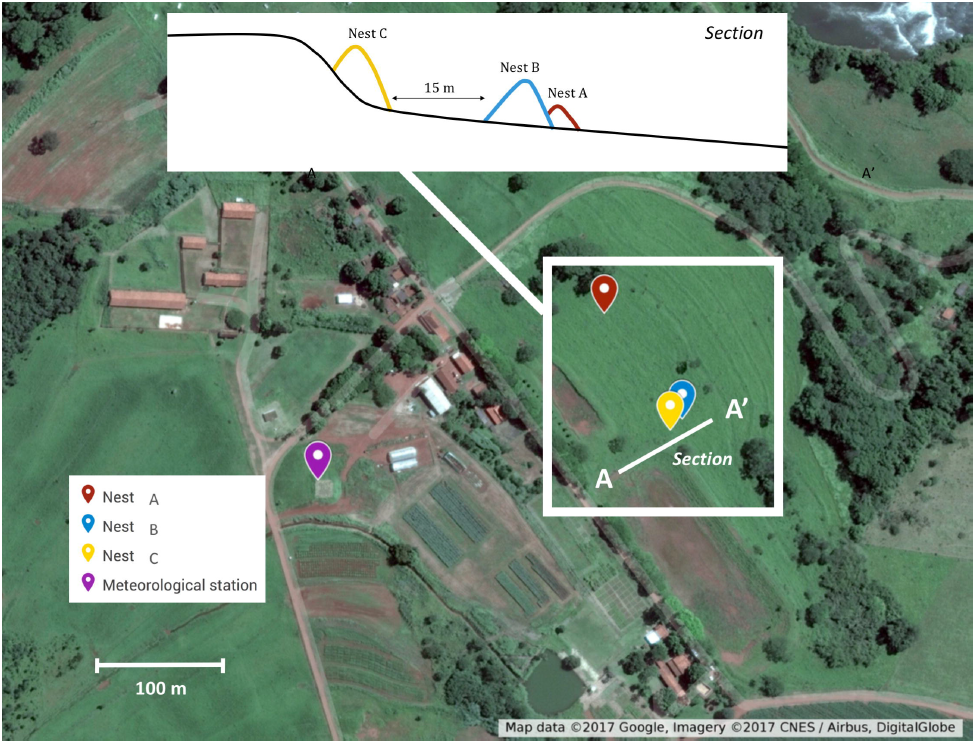
Experimental field area. The labels indicate the nest positions in the field. The line A/A’ indicates a sagittal section of the field (inset) showing the slope and altitude difference between the nest C and the two other nests (A and B). Map data ©2017 Google, Imagery © 2017, CNES/Airbus, DigitalGlobe.

**S2 Fig.**
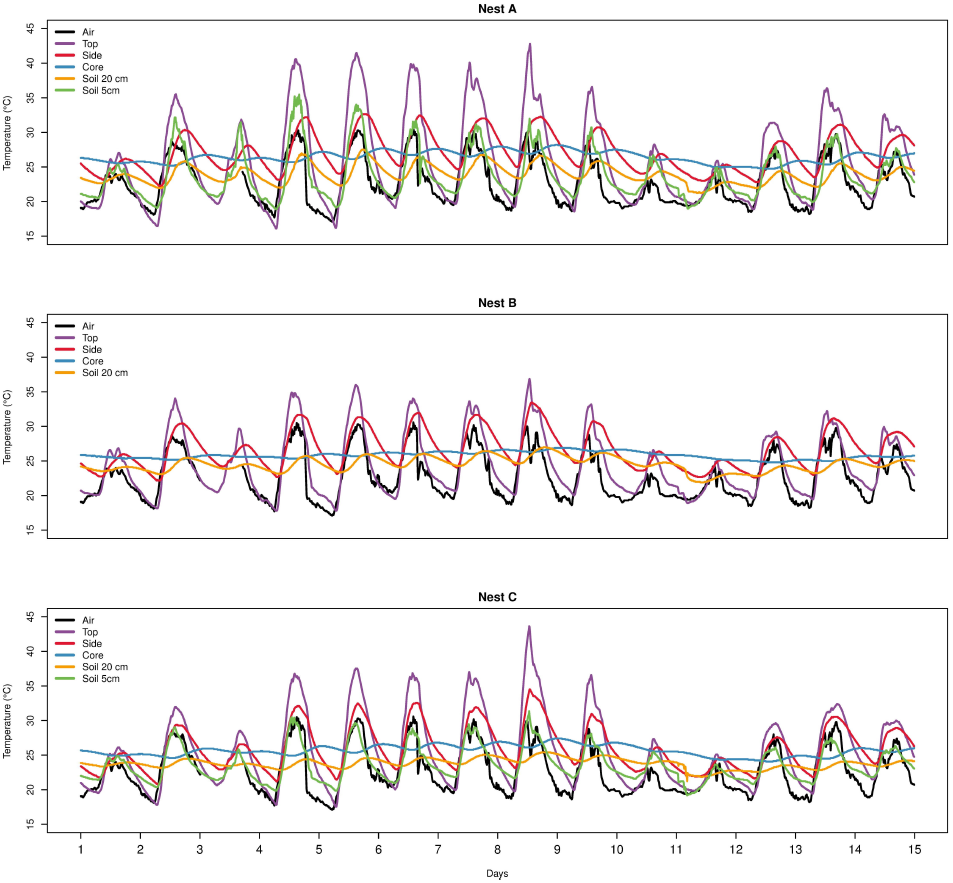
Raw time series data for the 3 nests. The lines are the raw temperatures as a function of time for the different probe locations.

